# MILDEW RESISTANCE LOCUS O (MLO) proteins function as trimeric inward calcium channels

**DOI:** 10.64898/2026.05.18.725938

**Authors:** David Ušák, Šárka Mattauchová, Michal Daněk, Roman Hudeček, George A. Caldarescu, Viktor Žárský, Roman Pleskot

**Affiliations:** Institute of Experimental Botany, Czech Academy of Sciences, Prague, Czech Republic; Department of Experimental Plant Biology, Faculty of Science, Charles University, Prague, Czech Republic; Department of Biochemistry and Microbiology, University of Chemistry and Technology, Prague, Czech Republic; Department of Comparative Development and Genetics, Max Planck Institute for Plant Breeding Research, Cologne, Germany

**Keywords:** Mildew resistance locus O, MLO, calcium channel, protein structure, molecular dynamics

## Abstract

Calcium signalling and structural roles are fundamental for plant growth, development, and environmental adaptation. Recent studies have identified MILDEW RESISTANCE LOCUS O (MLO) proteins as novel calcium-permeable channels with roles in root growth, cell wall development, pollen tube growth, and perception. However, the molecular mechanisms underlying MLO function remain unknown. Here, we demonstrate that multimerisation is essential for MLO activity. Chemical crosslinking, split-ubiquitin interaction assays, and single-molecule photobleaching revealed that MLO proteins form stable dimeric and trimeric assemblies at the plasma membrane. Structural modelling uncovered a molecular architecture of the MLO trimer with a central ion-conducting pore, which was further examined by molecular dynamics simulations in a lipid membrane environment. Computational electrophysiology showed preferential inward Ca^2+^ transport, confirming that MLO proteins function as calcium influx transporters, and identified a conserved set of pore-lining residues that coordinate ion translocation. Functional and structural analyses indicated that the mechanism of calcium permeation is evolutionarily conserved. Our findings provide mechanistic insight into MLO-mediated calcium influx across the plasma membrane and establish multimerisation as a critical determinant of this calcium channel’s activity.

## Introduction

Calcium is an essential macronutrient and a ubiquitous second messenger in plants, performing both structural and regulatory functions that are critical for growth, development, and environmental response^1^. As a structural component, calcium interacts with membrane negatively charged lipids^2^ or stabilises cell walls through cross-linking of pectins^3^. Along with this constitutive role, transient spatio-temporal changes in cytosolic Ca^2+^ concentration act as highly dynamic signalling events that encode information in response to external and internal stimuli, including drought, salinity, temperature fluctuations, pathogen attack, hormones, and mechanical stress^4–6^. These calcium signatures are decoded by specialised sensor proteins such as calmodulins or calcium-dependent protein kinases, which subsequently regulate downstream processes^4^. Because resting cytosolic Ca^2+^ levels must remain low while apoplastic calcium concentrations are high, tightly controlled transport across the plasma membrane is indispensable. Calcium-permeable channels mediate influx during signalling, whereas Ca^2+^-ATPases and Ca^2+^/H^+^ exchangers restore basal concentrations by extrusion or sequestration^6^. Thus, plasma membrane calcium transport, coordinated with tonoplast ion transport, is central not only to nutrient acquisition but also to the generation, propagation, and termination of calcium-based signals, which are essential for plant cell functions.

Several classes of calcium-permeable channels have been identified in plants, although their molecular diversity and regulation remain areas of active research. Among the best characterised are cyclic nucleotide-gated channels (CNGCs), which are non-selective cation channels activated or modulated by calmodulin and phosphorylation, and are implicated in pathogen defence, thermosensing, pollen tube growth, and development^7^. Glutamate receptor-like channels (GLRs), homologous to animal ionotropic glutamate receptors, mediate Ca^2+^ influx in response to amino acids and participate in long-distance wound signalling, root architecture, and stress responses^8^. Mechanosensitive channels, including members of the Mid1-Complementing Activity (MCA) family and OSCA/TMEM63 proteins, contribute to calcium entry triggered by touch, osmotic stress, or membrane tension^6,9,10^.

Recent studies have described MILDEW RESISTANCE LOCUS O (MLO) proteins as novel calcium-permeable channels with functions throughout the plant, including root growth, cell wall development, pollen tube growth and perception^11–13^. MLO proteins have been shown to function as calcium influx channels^14^, yet we lack the mechanistic understanding of how they transport calcium^15^. Here, we provide the first experimental evidence into the MLO oligomerisation, which leads to the formation of a pore at the subunit interface. Our computational data show that this pore is capable of inward translocation of Ca^2+^ ions, which is mediated by a set of evolutionarily conserved residues. Taken together, our work provides mechanistic insight into Ca^2+^ transport by MLOs and serves as a basis for future investigations into the complex regulation of calcium homeostasis in plants.

## Results and Discussion

Given that currently known plant calcium channels do not function as monomers^6^, we hypothesised that MLO proteins form higher-order assemblies to function as calcium channels. To assess the capacity of MLO proteins for mutual interaction, we employed a yeast split-ubiquitin assay (SUS). To evaluate the robustness of these interactions, we selected homologs from distinct Arabidopsis phylogenetic clades^16^. All tested combinations, including both homo- and heteromeric interactions, exhibited strong colony growth (**Fig. 1a**), indicating the formation of higher-order assemblies. To further examine MLO association *in planta*, we expressed MLO-mEGFP in *Nicotiana benthamiana* (**Fig. 1b**). All MLOs predominantly localised to the plasma membrane (**Fig. 1c**), a finding corroborated by subcellular fractionation (**Fig. 1d**), which supports previous reports that MLOs are primarily plasma membrane-resident proteins^17^. We then utilised MLO-enriched microsomal fractions (MF) from *N. benthamiana* as input for *in vitro* analyses of MLO oligomerisation. The MF fractions were incubated with bis(sulfosuccinimidyl)suberate (BS3), a bifunctional cross-linker that covalently links lysine residues of interacting proteins, thereby stabilising protein complexes^18^. Immunoblot analysis of the crosslinked MF fractions revealed the increased levels of higher-molecular-weight protein assemblies compared to the monomer (**Fig. 1e**). Relative to the ∼85 kDa monomer, the observed bands at 170 and 250 kDa corresponded to dimeric and trimeric assemblies of MLO, respectively. Additionally, we conducted a single-molecule photobleaching experiment^19^ to independently determine the stoichiometry of MLO complexes in the isolated non-crosslinked MF fraction. Consistent with our crosslinking results, we observed that MLO proteins exist as monomers, dimers, and trimers, further supporting the notion that the MLO oligomerisation occurs naturally in plant cells (**Fig. 1f)**.

**Figure 1:**
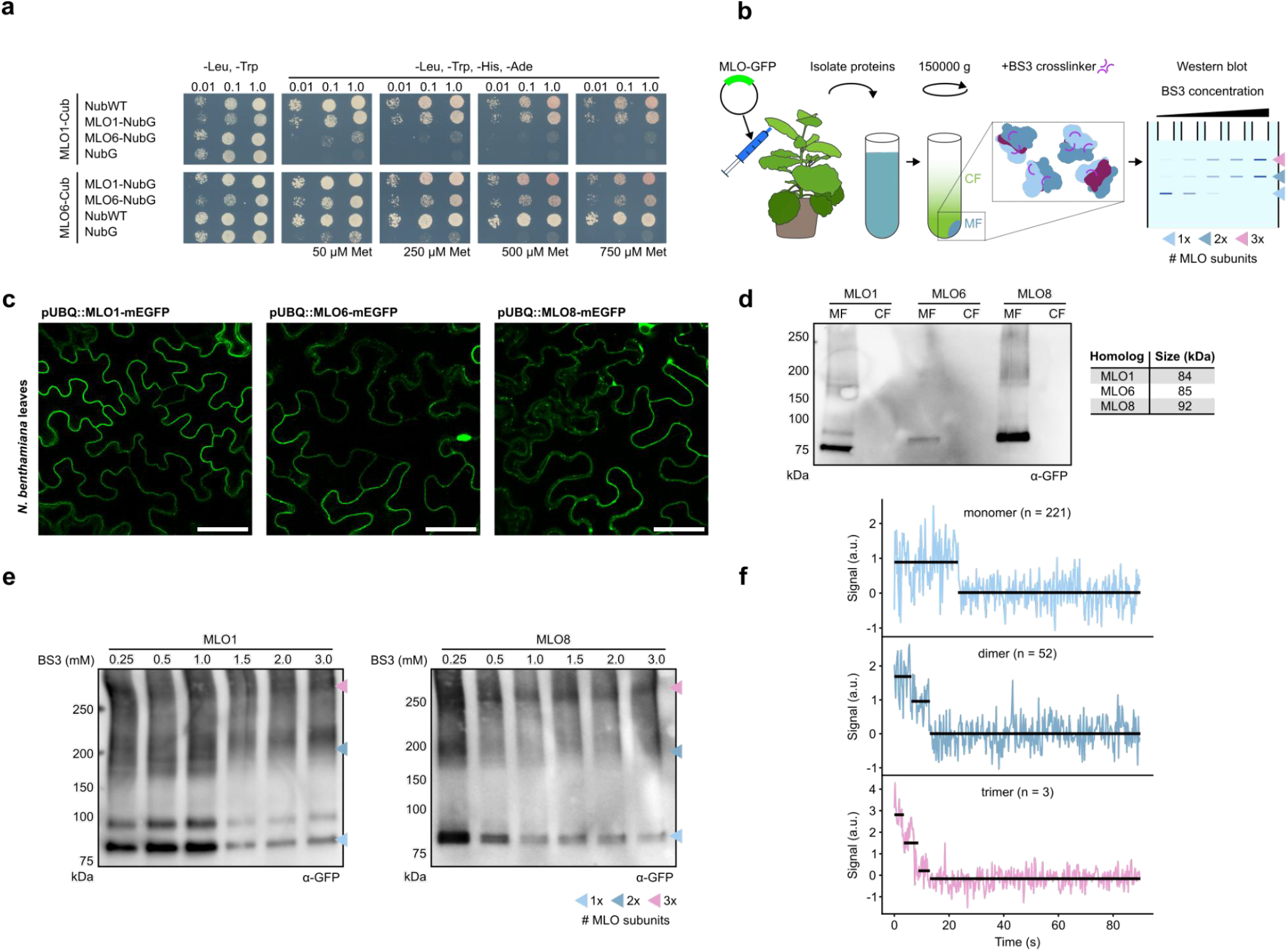
MLO proteins assemble into dimers and trimers *in vivo*. (**a**) Split-ubiquiting assay of MLO1 and MLO6. The Cub or Nub-tagged MLO homologs were coexpressed in yeast, with the colony growth reflecting the ability of specific combinations to interact. NubWT and NubG were used as positive and negative controls, respectively. (**b**) Diagram of the experimental approach for the MLO oligomerisation analysis *in planta*. The MLO-GFP cassette is transiently expressed in *Nicotiana benthamiana*. After 3 days, the proteins from leaves are isolated and fractionated to the microsomal fraction (MF, pellet) and the cytosolic fraction (CF, supernatant). The protein complexes in the MF are stabilized by a crosslinker, with the stoichiometry analyzed by SDS-PAGE and Western blot. (**c**) Imaging of MLO-mEGFP in *Nicotiana benthamiana* leaves. The proteins are predominantly localized to the plasma membrane. Scale bar = 50 µm. (**d**) Subcellular fractionation of protein isolates from (**c**). The enrichment of MLO-mEGFP in the microsomal fraction (MF) compared to its absence in cytosolic fraction (CF) validates the membrane residence of MLOs. (**e**) Crosslinking of the MLO1 and MLO8 MF fraction by bis(sulfosuccinimidyl)suberate (BS3) and subsequent analysis by immunoblotting. With the increased BS3 concentration, the higher-size bands corresponding to dimeric and trimeric stoichiometry (dark blue and burgundy arrows) are enriched, while the monomer band (light blue) is reduced. Load = 5 µg of the MF fraction. (**f**) Single-molecule photobleaching of MLO8-mEGFP. Representative fluorescence traces of monomeric, dimeric, and trimeric assemblies, respectively. Each trace shows the background-subtracted fluorescence intensity within a 5 × 5 pixel region of interest under continuous 488 nm illumination. Black horizontal lines indicate the mean fluorescence intensity of each plateau between consecutive bleaching events. a.u. = arbitrary units.

The SUS, crosslinking results, and single-molecule photobleaching data imply that MLOs interact to form higher-order complexes comprising two or three subunits. To analyse the molecular architecture of these assemblies, we used AlphaFold2-Multimer 3 (AF2-MM3) to predict MLO1, MLO6, and MLO8 dimers and trimers. AF2 prediction of MLOs yielded highly-confident models for all homologs, consisting of multiple transmembrane (TM) helices folded into a well-defined structure and a calmodulin-binding domain, connected by an unstructured region^15^ (**Fig. S1a**,**b**). Next, given the highly similar structure of all investigated MLOs, we selected MLO6 as a representative isoform. Within the MLO6 dimer, the TM regions interact to form a single integral membrane region and two CAM-binding domains are positioned towards the cytoplasm (**Fig. 2a**). The MLO6 trimer adopts a similar topology, with the CAM-binding domains buried slightly within the emerging cytoplasmic cavity. Compared to the dimer, the subunits rotate approximately 40° away from each other to facilitate the introduction of the third subunit (**Fig. 2a**). This leads to the formation of a triangular shape that creates a small pore at the central interface of the three subunits (**Fig. 2b**). The overall quality of the obtained complexes is validated by the AF2 global confidence metric (**Fig. 2c, Fig. S1b**). Moreover, the mapping of the structural features to the pLDDT score revealed that the trimer interface residues lie within the highly-scoring regions, further supporting the predicted model (**Fig. 2d**). In a likewise manner, we also predicted MLO6 oligomers of up to 6 subunits. However, in the 4-6 subunit predictions, the ipTM score dropped sharply, suggesting that the trimer is indeed the largest relevant MLO complex (**Fig. 2e**).

**Figure 2:**
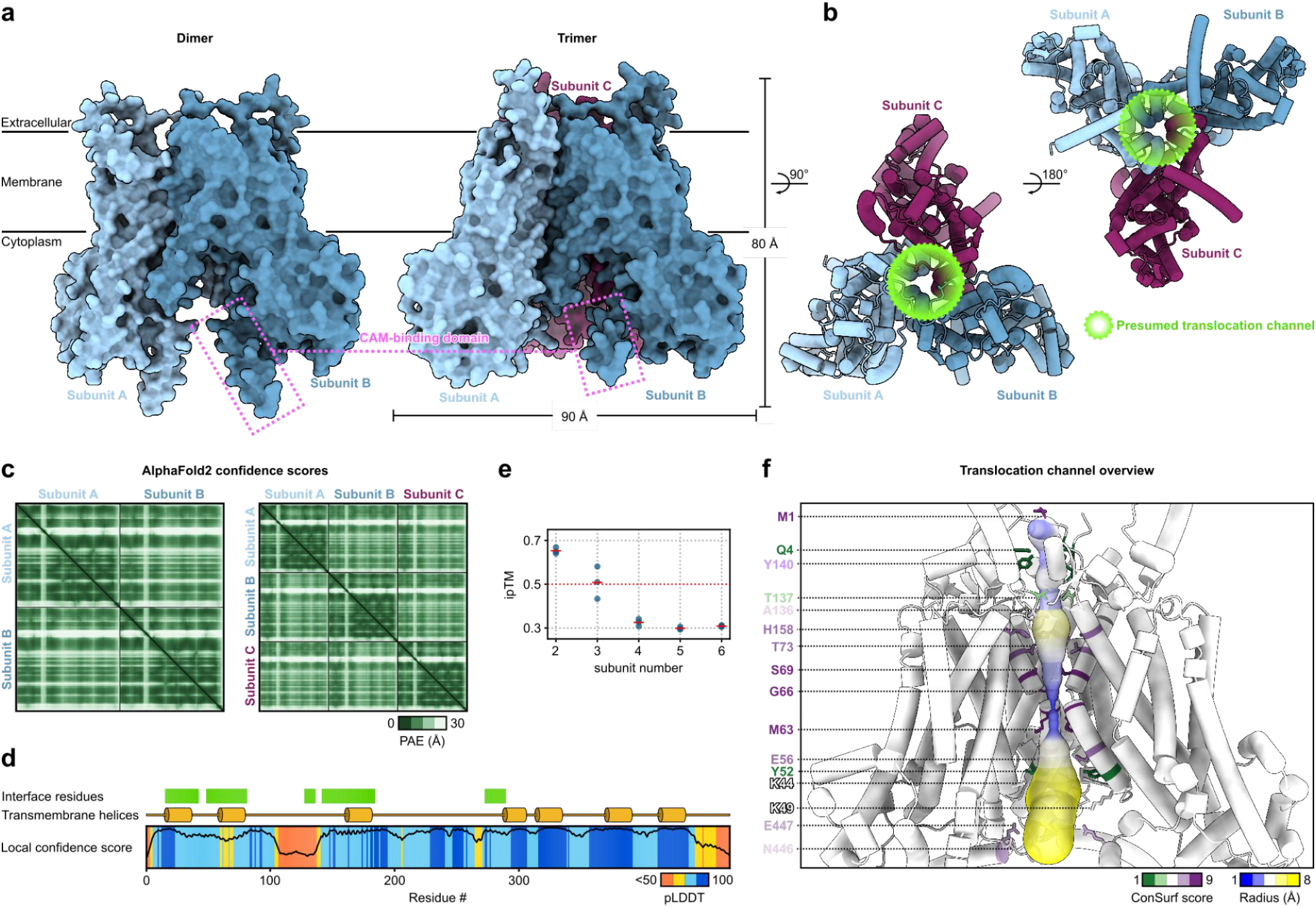
The MLO trimer forms a pore across the lipid bilayer. (**a**) AlphaFold2 predictions of MLO6 as a dimeric or a trimeric assembly. The surface representation is shown. The position of the assembly in relation to the membrane is depicted (**b**) Top and bottom view of the MLO6 trimer in the tube representation. The individual subunits adopt a triangular shape, with a possible translocation channel forming at the centre of the interface.(**c**) AlphaFold2 confidence for the MLO6 oligomeric assemblies. The predicted alignment error (PAE) plot shows high confidence of both dimer and trimer predictions. (**d**) AlphaFold2 confidence, the pLDDT score, for the single MLO6 chain. The position of the trimer interface residues and transmembrane helices are highlighted. These structural features are located at the highly-confident regions (**e**) Overall model confidence for the MLO6 oligomers consisting of 2-6 subunits. Only dimers and trimers exhibit a high ipTM score. (**f**) The MLO6 trimer translocation channel as determined by the MOLE algorithm. The channel is shown in the balloon representation, coloured by its radius. The pore-lining residues, coloured by ConSurf, are indicated to the left.

The presence of an opening at the centre of the trimer assembly supports the hypothesis that MLO oligomerisation enables their channel-like properties^15^. To identify putative pores that promote calcium transport through the structure, we used the MOLE software to predict a possible pore^20^ in the MLO6 trimer. The algorithm identified a single pathway of approximately 139 Å in length, traversing from the extracellular side to the cytoplasm. The resulting pore is formed at the central interface of the three subunits, whose TM2 helices create a 1 Å bottleneck in the middle of its trajectory (**Fig. 2f**). Furthermore, we subjected the MLO6 trimer to a ConSurf analysis of protein evolutionary conservation^21^. Upon mapping to the pore-lining residues, we observed multiple highly conserved residues, oriented towards the central cavity. Among these, the M63, G66 and S69 exhibited the highest score, with E56, T73, A136, Y140, H158, N446 and E447 all showing an above-average conservation (**Fig. 2f**).

The high evolutionary conservation of protein-protein interface residues, such as at the MLO6 trimer pore bottleneck, likely underlies their functional importance^22^. To study the ability of the MLO trimer structure to transport calcium ions, and to analyse the mechanical role of the interface residues during the translocation, we employed all-atom molecular dynamics (MD) simulations of the MLO6 homotrimer. As such, the protein was inserted into a lipid bilayer and the system included 50 mM CaCl_2_. After 500 ns of simulation time, no transport of Ca^2+^ across the membrane occurred. However, under these conditions we noticed a slight relaxation of the subunits, leading to an enlargement of the central pore, with the bottleneck extending to 2.5 Å (**Fig. 3a**). Therefore for our subsequent simulations, we utilised a 100 ns relaxed trimer as an input structure (**Fig. 3b**). Moreover, we applied an external electric potential of either +700 or -700 mV to the system, promoting the outward or inward translocation, respectively (**Fig. 3c**). Similar values were used in previous studies, and it is generally accepted as an approximation of conditions present in living cells^23,24^. After 500 ns, we observed inward transport of multiple Ca^2+^ ions in all three simulation repeats, while only one Ca^2+^ ion was transported in the outward direction in one simulation replica (**Fig. 3d**,**h; Fig. S2**). While these results support the MLO trimer as a Ca^2+^-translocating protein, the application of external electric potential, with a need to restrain a protein structure to avoid protein destabilisation, does not fully reproduce the biological processes driving channel-mediated transport. Thus, we adopted the computational electrophysiology approach, which uses ion concentration gradients to generate the electric potential, more closely reflecting the natural conditions and allowing us to study the protein dynamics without restraining it^25^. In this case, the simulation system contained the relaxed MLO6 homotrimers within two lipid bilayers, on top of each other, together forming an inner and outer compartment. To account for the transport directionality, each trimer was placed in a mirroring orientation in relation to the central compartment. A concentration of 50 mM CaCl_2_ was used to provide the transported ions (**Fig. 3e**). A mismatch of 12 Ca^2+^ ions was maintained between the compartments, corresponding to approximately 900 mV transmembrane potential (**Fig. 3f**). The simulation was computed for 500 ns simulation time, after which we recorded Ca^2+^ transport exclusively for the inward direction. Although some outward-bound ions entered the cytoplasm-adjacent cavity of the trimer, they were not able to translocate to the next compartment (**Fig. 3g**). After increasing the Ca^2+^ mismatch to 20 molecules (or 1250 mV), this directional preference was even more pronounced, with 18 inward translocations and only one ion passing in the outward direction (**Fig. 3h**). Altogether, these results demonstrate that the MLO6 homotrimer is well capable of mediating Ca^2+^ influx, and thus act as an inward membrane channel.

**Figure 3:**
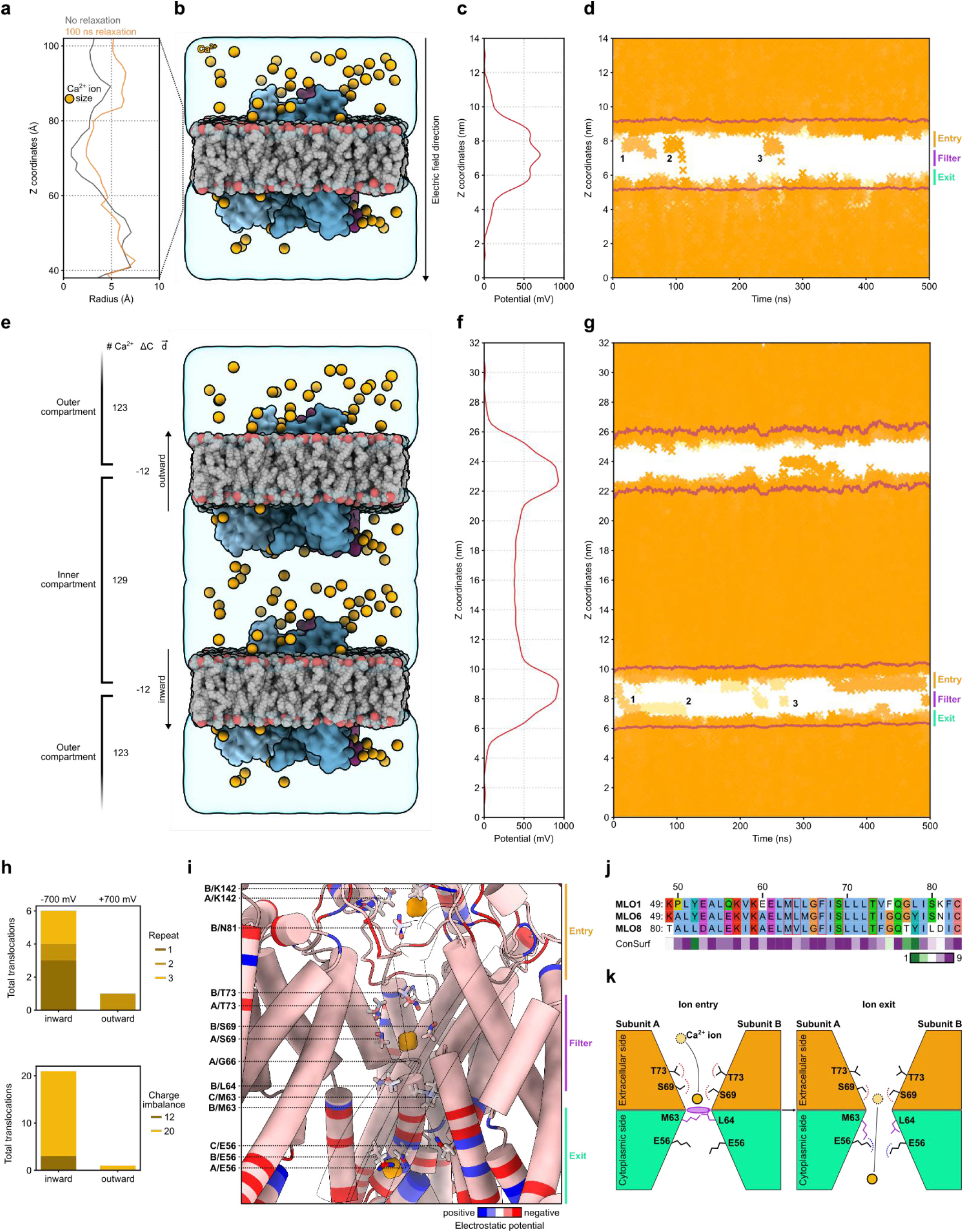
The MLO6 trimer is a calcium influx channel. (**a**) Radius of the non-relaxed structure compared to the MLO trimer equilibrated for 100 ns. The relaxation enables opening of the pore necessary to accommodate the Ca^2+^ ion. (**b**) Overview of the single membrane simulation setup. The protein is embedded in a POPC/POPE membrane. For the production run, an external electric potential is applied to the system, promoting the transport of Ca^2+^ ions. (**c**) Spatial distribution of the electric potential applied during the simulation. (**d**) Z axis trajectories of the Ca^2+^ ions in time. The individual translocation events are marked by numbers. (**e**) Overview of the double membrane system. The proteins are embedded in two stacked membranes, forming an isolated compartment in the center. The parameters of the outer and inner compartments are described to the left. (**f**) Spatial distribution of the electric potential induced by the ion mismatch between the two compartments. (**g**) Z axis trajectories of the Ca^2+^ ions in time. The individual inward translocation events are marked by numbers.(**h**) Quantification of the MLO6-dependent Ca^2+^ transport. The transport preferentially occurs in the inward direction. (**i**) Translocation states of Ca^2+^ and the pore-lining residues. Each Ca^2+^ atom (orange) represents a different step of the transport into the cell. The interacting residues are coloured by their electrostatic potential. (**j**) Multiple sequence alignment of MLO1, MLO8 and MLO6 central pore residues. The residues active in ion translocation are highlighted in red. The evolutionary conservation of each residue is shown by the ConSurf score below. (**k**) Presumed MLO6-dependent transport mechanism of Ca^2+^. Upon entry to the tunnel, the ion is lured by polar residues above the selectivity filter. The selectivity filter sterically inhibits further transport. After the movement of H63 and L64 away from the filter, the Ca^2+^ ions are transported further and eventually exit the pore.

Next, we investigated the molecular mechanisms of the calcium transport in detail. The MD trajectories revealed that during the inward translocation event, the ion first enters the channel from the extracellular side, followed by a prolonged retention above the centre of the complex and subsequent quick expulsion to the cytoplasm (**Fig. 3d**,**g**). Based on the Ca^2+^ position during the import, we marked these events as Entry, Filter and Exit, respectively. Subsequently, we analysed the pore-resident amino acids and their interaction with the Ca^2+^ ion during these translocation steps. At the Ca^2+^ entry, the ion is coordinated by the side chains of K142 and N81 (**Fig. 3i**). These residues are located in the MLO6 extracellular loop, which possibly acts as an entrance plug, providing preliminary selection of the transported ion. After entering the channel, the ion interacts with S69 and T73 (**Fig. 3i**). Hydroxyl groups of these amino acid residues coordinate the positively charged Ca^2+^ and guide it towards the bottleneck at the centre of the MLO6 interactory interface, acting as a selectivity filter. The selectivity filter itself is formed by a short loop containing the highly conserved G66, which leads to a kink in the TM2 structure. Directly underneath the kink, the side chains of M63 and L64 act together, preventing the ion movement (**Fig. 3i**). Their rotation away from the bottleneck provides space for Ca^2+^ passage into the Exit position, where the Ca^2+^ ion is attracted by the negatively charged E56 (**Fig. 3i**). At this position, the tunnel enlarges, feeding the translocated ion into the cytoplasm. Interestingly, at the very end of the Exit site, a lysine-rich motif is located (**Fig. 3i**), its high positive charge likely facilitating the Ca^2+^ expulsion.

While the MD simulation of the MLO6 trimer provided a detailed description of the mechanism of calcium transport by MLOs, it provided only limited information about functional conservation in other homologs. We therefore also examined the ConSurf score of the Ca^2+^-coordinating regions, particularly the TM2. Indeed, the residues shown to directly interact with the translocated ion are conserved in both MLO1 and MLO8 (**Fig. 3j**). Moreover, the residues E56, M63, L64, S69 and T73 exhibit almost full conservation also in other species, substantiating their importance^16^ (**Supplementary data**). Based on these data, we propose a general model of calcium import by the MLO trimer complex (**Fig. 3k**).

Taken together, our experimental results and computational simulations strongly support the hypothesis that MLO oligomerisation mediates calcium transport through the PM. As such, this work validates MLO as a novel player in the maintenance of plant calcium homeostasis.

## Materials and Methods

### Cloning and plant transformation

The coding sequences of *Arabidopsis thaliana* L. Heynh MLO1 (AT4G02600), MLO6 (AT1G61560) and MLO8 (AT2G17480) were domesticated into the GoldenGate-compatible entry vector pUPD2-CD. The MLO cassette was then recombined with pGGA-UBQ10-B, pGGB-linker-C, pGGD-mEGFP-E, pGGE-tHSP18-F and pGGF-linker-G into the pGGK destination vector^26^. The expression vectors were introduced into the GV2260 *Agrobacterium tumefaciens* strain and *Nicotiana benthamiana* leaves were infiltrated according to the original protocol^27^. MLO1 and MLO6 coding sequences were introduced between SalI and NotI restriction sites in pENTR1a Gateway compatible plasmid (Thermo Scientific) by restriction/ligation cloning. Subsequently, expression plasmids suitable for SUS were generated using Gateway LR Clonase II (Thermo Scientific) recombination from pMetYC-Dest and pXN22-Dest^28^.

### Confocal microscopy

Transformed *N. benthamiana* leaf epidermal cells were imaged three days after infiltration. 5 mm leaf disk was cut out from the leaf abaxial side, transferred to a water drop on a glass slide and observed on a Zeiss LSM880 confocal microscope with C-Apochromat 40x/1.2 W objective. The GFP was excited using a 488 nm laser at 2.0% intensity, the emission was recorded at 493-574 nm. The same imaging setup was used for all samples.

### Protein isolation and subcellular fractionation

*N. benthamiana* leaves were cut off from the plant and frozen in liquid nitrogen. For the protein isolation, the frozen leaves were ground to a fine powder in a mortar filled with liquid nitrogen. The material was then homogenised further using Homogenisation buffer (50 mM HEPES, 400 mM sucrose, 100 mM KCl, 100 mM MgCl_2_, pH 7.5) and then filtered through a nylon mesh. To remove the cell debris, the extract was centrifuged at 15,000 g (15 min, 4°C) twice, with the supernatant transferred to a new tube in between. Subsequently, the extract was fractionated at 150,000 g (60 min, 4°C), separating the cytosolic fraction (CF, supernatant) from the microsomal fraction (MF, pellet). The obtained pellet was then resuspended in phosphate buffer (5 mM KH_2_PO_4_, 5 mM Na_2_HPO_4_, pH 7.8) and kept on ice for future handling.

### Protein crosslinking

Microsomal fractions were incubated with varying concentrations of BS3 (Thermo Fisher) and crosslinked on a vertical rotator for 2 hours at 4 °C. After crosslinking, the reaction was quenched by adding glycine to a final concentration of 50 mM and incubated for 15 min at room temperature. Crosslinked samples were analysed by Western blotting.

### SDS-PAGE, Western blot and immunodetection

Protein samples prepared by the aforementioned biochemical approaches were mixed with Laemmli buffer and XT reducing agent (both Bio-Rad), boiled at 55°C for 10 mins and separated on 4-20% stain-free polyacrylamide gels (Bio-Rad) at 180 V for 60 min. Next, the proteins were transferred to a Trans-Blot Turbo PVDF membrane (Bio-Rad) at 12 V for 10 min. Membranes were rinsed three times in 1xTBS-T buffer, followed by overnight blocking in 5% non-fat milk in TBS-T. The membrane was incubated with mouse α-GFP (Sigma) and HRP-conjugated α-mouse (Promega) antibodies, with TBS-T washes in between. Afterwards, the HRP-derived chemiluminescent signal was detected using Radiance Q substrate (Azure) with ChemiDoc Imaging System (Bio-Rad).

### Single-molecule photobleaching analysis

MF containing MLO8–mEGFP were loaded onto cleaned glass coverslip. Fluorescent time-lapse images were acquired on a spinning-disk confocal microscope (camera directly coupled to the spinning-disk unit, without a microscope body) equipped with a 100× / 1.4 NA oil-immersion objective, yielding a pixel size of 0.11 µm. Stacks of 360 frames were recorded under continuous 488 nm laser illumination (25% power), with an exposure time of 250 ms per frame.

Maximum-intensity projection was generated from the full image stack. Spots were detected using the ComDet plugin in FIJI (v0.5.6)^29^ with intensity threshold = 3, particle size parameters = 3 pixels. To exclude noise detections and macromolecular clusters, spots were filtered by size and shape in the next step. The custom Python script extracted per-spot fluorescence traces from the raw image stacks. For each detected spot, fluorescence intensity was computed as the mean of a 5 × 5 pixel region of interest (0.55 × 0.55 µm) centered on the spot. Local background was estimated for each frame as the mean intensity of an annular region (inner radius 4 px / 0.44 µm; outer radius 7 px / 0.77 µm) surrounding the region of interest (ROI), and subtracted from the signal. Spots within seven pixels of the image edge were excluded. The background-subtracted traces were inspected using a custom interactive Python tool. Traces were presented in randomised order to avoid selection bias. For each trace, bleaching transitions were marked manually; traces without resolvable steps were marked as unusable and excluded. Step amplitudes were computed as the difference between the mean intensities of the segments before and after each marked transition. For each scored trace, the total bleach amplitude (mean intensity over the first 20 frames minus the mean over the final 50 frames) was expressed in units of the unitary step (bleach_units). Traces with |bleach_units - n_steps| > 0.5 were excluded as inconsistent between scored step count and total bleach amplitude, accounting for both over-segmentation (noise scored as steps) and missed steps (insufficient intensity drop for the scored count). Out of all manually scored spot traces (1253), 276 passed quality filtering. The distribution of step counts was tested against truncated binomial models (truncated because molecules with all dark fluorophores cannot be observed) for candidate stoichiometries n = 2, 3, and 4.

### Split-ubiquitin system

The THY.AP4 yeast strain was cotransformed with pMetYC (bait) and pXN22 (prey) vectors armed with the MLO isoforms under investigation by applying the lithium acetate/single-stranded carrier DNA/PEG method^28^ and plated on YNB + CSM (both MP Biomedicals) medium lacking Leu and Trp supplemented with 2% glucose and 50 μM Met. After a 2-day recovery at 30°C, freshly grown colonies were resuspended in milliQ water and diluted to obtain suspensions of optical densities (OD600) of 1.0, 0.1, and 0.01. Yeast suspensions of 8 μl were dropped onto plates with YNB + CSM selective medium lacking Leu, Trp, Ade, and His, supplemented with 2% glucose and 50, 250, 500 and 750 μM Met. Yeast growth was visually assessed after incubation for 2-3 days at 30°C. The non-recombined pNX35 vector encoding for NubG unable to reassemble with Cub, and wild-type Nub moiety spontaneously reassembling with Cub, were used as negative or positive control for each bait, respectively.

### Protein structure prediction, analysis and molecular dynamics simulations

To generate the structural models of oligomeric MLOs, AlphaFold2-MM3 was employed^30^. For all predictions, we used the full length sequence of the protein. The overall quality and confidence of the model was determined from the pLDDT and ipTM scores, as well as the PAE plots.

To identify the putative translocation site, the obtained structure was submitted to MOLE online tool^20^, set at pore mode (default settings). The longest and the best scoring pore was subsequently selected.

ConSurf analysis of evolutionarily conserved residues was employed to map the evolutionary variability of the MLO structure^21^. The default settings were used for all analyses.

All molecular dynamics (MD) simulations were executed using the GROMACS software version 2020^31^. The atomic representation AF2 structures were converted to the CHARMM36m forcefield^32^ in the CHARMM-GUI v3.8 web suite^31^. Briefly, the simulated MLO6 trimer was embedded in a lipid bilayer (POPC and POPE ratio 1:1). Multisite calcium atoms were added to the system as the translocated substrate^33^. To simulate the transmembrane gradient, an external electric potential of +/-700 V was applied; to prevent undesirable structural deformations, the protein backbone was fixed by harmonic restraints of 1000 kJ.mol^-1^.nm^-2^. Alternatively, we also utilised the GROMACS electrophysiology framework^25^. All simulations were run for 500 ns, after which the individual trajectories were investigated into detail, particularly the MLO channel-mediated Ca^2+^ permeations. All simulation parameters for MD runs are deposited in the Zenodo repository.

## Data availability

Custom scripts for single-molecule photobleaching experiments are available in the GitHub repository. Computational results and structure-related data files are deposited in the Zenodo repository.

## Acknowledgment

This work was supported by Czech Science Foundation grant nr. 22-35680M (R.P.). Microscopy was performed at the Imaging Facility of the IEB CAS, supported by MEYS CR LM2023050 ‘Czech-BioImaging’ and IEB CAS. We acknowledge VSB – Technical University of Ostrava, IT4Innovations National Supercomputing Center, Czech Republic, for awarding this project access to the LUMI supercomputer, owned by the EuroHPC Joint Undertaking, hosted by CSC (Finland) and the LUMI consortium through the Ministry of Education, Youth and Sports of the Czech Republic through the e-INFRA CZ (grant ID: 90254).

---

**Figure S1:**
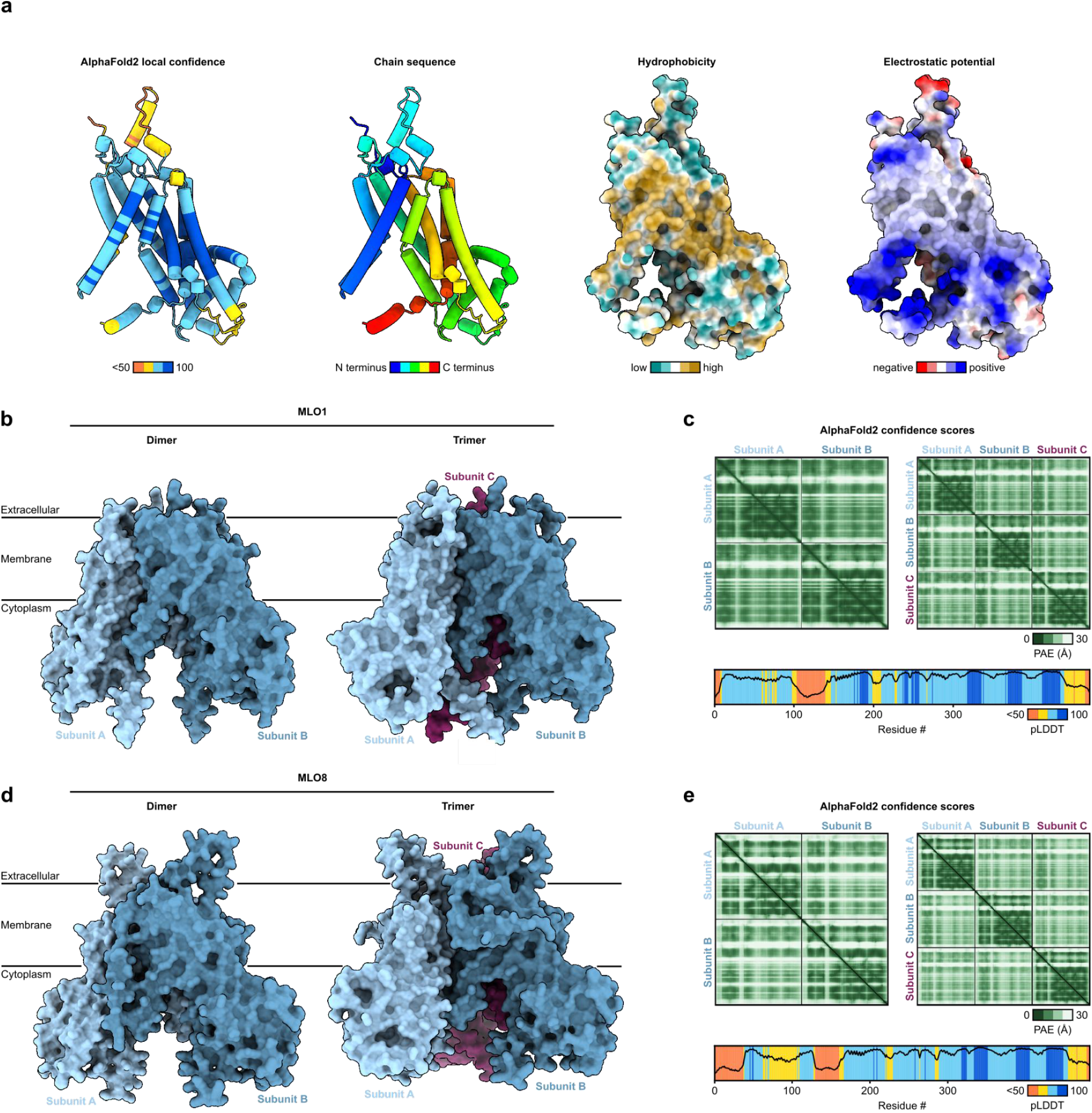
AlphaFold2 (AF2) prediction of MLO proteins. (**a**) MLO6 adopts a well folded helical structure. High pLDDT metric supports the model confidence. Chain order coloring shows the topology of the helices constituting the structure. The transmembrane helices assemble into a single integral membrane region, as revealed by residue hydrophobicity. Coulombic electrostatic potential shows multiple charged hotspots on the protein surface. (**b**) AF2-MM3 predictions of MLO1 dimer and trimer. (**c**) Confidence metrics for the prediction in **b.** (**d**) AF2-MM3 predictions of MLO8 dimer and trimer. (**c**) Confidence metrics for the prediction in **d**.

**Figure S2:**
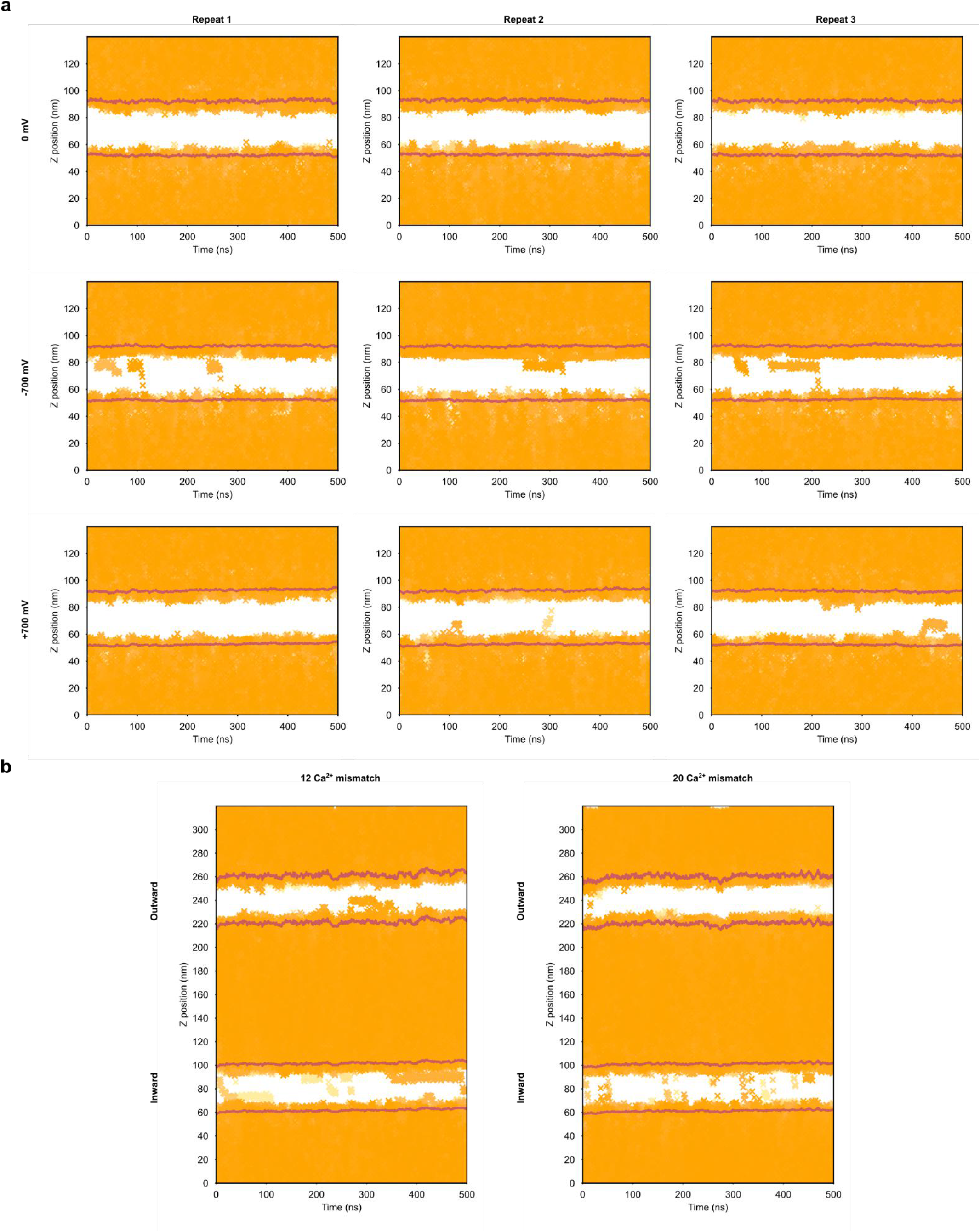
Computational electrophysiology of MLO6-mediated Ca^2+^ transport. (**a**) Simulation repeats of a single-membrane system containing the MLO6 trimer with no (top), negative (centre) or positive (bottom) transmembrane potential applied. (**b**) Double membrane system with transmembrane potential simulated by 12 (left) and 20 (right) molecule Ca^2+^ mismatch between the outer and inner compartments. The translocations in **a** and **b** can be observed preferentially in the inward direction.

